# The abiotic and biotic environment together predict plant, mammal, and bird diversity and turnover across the United States

**DOI:** 10.1101/2021.11.11.468263

**Authors:** Amanda S. Gallinat, William D. Pearse

## Abstract

The distribution of taxonomic, phylogenetic, and functional biodiversity results from a combination of abiotic and biotic drivers which are scale dependent. Parsing the relative influence of these drivers is critical to understanding the processes underlying species assembly and generating predictions of biodiversity across taxonomic groups and for novel sites. However, doing so requires data that capture a spatial extent large enough to reflect broad-scale dynamics such as speciation and biogeography, and a spatial grain fine enough to detect local-scale dynamics like environmental filtering and biotic interactions. We used species inventories of vascular plants, birds, and mammals collected by the U.S. National Ecological Observatory Network (NEON) at 38 terrestrial field sites, to explore the processes underlying taxonomic, phylogenetic, and functional diversity and turnover. We found that, for both species richness (alpha-diversity) and turnover (beta-diversity), taxonomic, phylogenetic, and functional diversity are weak proxies for one-another, and thus may capture different species assembly processes. All diversity metrics were best predicted by a combination of abiotic and biotic variables. Taxonomic and phylogenetic richness tended to be higher at warmer, wetter sites, reflecting the role energy inputs play in driving broad-scale diversity. However, plant diversity was negatively correlated with bird phylogenetic and mammal functional diversity, implying trait conservation in plant communities may limit niche availability for consumer species. Equally, turnover in bird and mammal species across sites were associated with plant turnover. That the biodiversity of one taxon is predictive of another across these North American sites, even when controlling for environment, supports a role for the cross-clade biotic environment in driving species assembly.

## Introduction

Patterns in the distribution of taxonomic, phylogenetic, and functional biodiversity across taxa can inform our understanding of the processes underlying community assembly (Cavender-Bares et al. 2009, Jetz et al. 2012a). In particular, parsing the relative contributions of abiotic (environmental) and biotic (other species) factors to community assembly and biodiversity is critical to predicting the direct and indirect impacts of environmental change and species loss in plant and animal communities. Abiotic and biotic drivers operate and interact across spatial scales (Westgate et al. 2014, Fine 2015). Classically, biogeography and speciation are considered broad-scale drivers while environmental filtering and interspecific interactions are thought to dominate local-scale assembly (Whittaker et al. 2001, Wang et al. 2009, Belmaker and Jetz 2011). Understanding how these broad- and local-scale drivers work in tandem to influence site-level biodiversity is an important next step for linking academic work on macro-ecological patterns with practical conservation decision-making at local sites. The relative contributions of the processes generating and maintaining biodiversity can be further informed by comparing patterns in richness (alpha-diversity) and turnover (beta-diversity) across assemblages. Turnover between sites can provide valuable context for broad patterns of alpha-diversity, reflecting heterogeneity in the taxonomic, phylogenetic, and functional identities of different assemblages. Furthermore, prior studies show taxonomic, phylogenetic, and functional diversity are often unreliable proxies for one another (Devictor et al. 2010, Cadotte and Tucker 2018), and understanding how and where they differ can provide insight into ecological and evolutionary processes. Parsing the drivers of each dimension of biodiversity independently presents an opportunity to examine, from multiple vantage points, how and why assemblages vary across biotic and abiotic gradients.

Investigations into what drives taxonomic, phylogenetic, and functional richness across sites have shown strong abiotic controls, particularly at the regional and global scale. Latitude and elevation are often strong regional predictors of biodiversity, and greater energy inputs (*e.g*., temperature, primary production) may lead to higher alpha-diversity across taxa (the ‘energy hypothesis’; Currie et al. 2004, Tittensor et al. 2010). The proposed mechanisms underlying this relationship include direct environmental (abiotic) impacts on speciation rates, as well as biotic drivers of speciation and niche availability (*e.g*., producer diversity driving consumer diversity; Hawkins et al. 2003, Belmaker and Jetz 2011). Efforts to parse these abiotic and biotic drivers of richness—and to understand whether cross-taxon congruence is driven by shared environmental responses or causal biotic links—have produced conflicting results across scales and taxa (Westgate et al. 2014). Studies covering sufficiently broad geographic extents to capture the impacts of biogeography and speciation on biodiversity suggest that once environmental responses are accounted for, the biodiversity of other taxa provide little additional predictive power (Jetz et al. 2008). However, those covering smaller geographic and environmental extents have found links among taxa beyond shared environmental responses (Kissling et al. 2007, 2008, Barrio et al. 2016).

While the patterns and drivers of total local biodiversity, or richness, can broadly indicate how biodiversity is generated and maintained, they may differ from the patterns and drivers of species composition, as represented by biodiversity turnover. The focus of turnover on differences in species identities across sites suggests similar rates of turnover among taxa could result from links related to resource use, interactions, and shared microhabitat responses not detected by broad-scale models. Compared to biodiversity richness, however, the processes underlying turnover are more poorly understood (Swenson et al. 2012), despite the importance of turnover in disentangling the mechanisms underlying biodiversity richness. For instance, the homogenization of diversity (*e.g*., in urban ecosystems) can result in high richness but low turnover (Groffman et al. 2014). Understanding regional-scale turnover is therefore important in and of itself for understanding what processes contribute to community composition, and it is also important for addressing ongoing debates about the drivers of alpha diversity (Gonzalez et al. 2016). In some cases, the diversity of other co-occurring taxa has been shown to be a stronger predictor of turnover than climate (Buckley and Jetz 2008), but the patterns and drivers of turnover have been shown to be highly scale-dependent (Barton et al. 2013, Mori et al. 2018).

Efforts to parse the drivers of both richness and turnover have ultimately been limited by a mismatch of scale. Data limitations often limit studies of broad spatial extents to aggregate data and so sacrifice fine-scale geographic resolution. A common concern in such studies of species whose ranges overlap is whether they can capture true co-occurrences or direct biotic interactions. Conversely, studies that do capture such fine-grained interactions do so across limited spatial extents, making it is difficult to capture broad environmental variation or the influence of biogeography. These practical constraints have made it challenging to explore the combined influence of abiotic and biotic processes on local-scale biodiversity. To overcome this challenge, what is needed are fine-grain, cross-taxon species assemblage data that cover a broad spatial extent.

The US National Ecological Observatory Network (NEON) is now emerging as a resource that offers fine grain, cross-taxon species inventories across the United States. These data provide a unique opportunity to investigate the contributions of the abiotic and biotic environment to site-level biodiversity. Here we explore NEON’s observations of vascular plants, birds, and small mammals to examine the patterns and drivers of alpha- and beta-diversity across the US. We examine taxonomic, phylogenetic, and functional diversity to generate insights into the patterns and drivers of species richness, biogeography, and niche differences, as well as the extent to which these processes reflect and interact with one another. Leveraging NEON’s fine spatial grain and broad spatial extent, we parse the abiotic and biotic predictors of biodiversity. We find that for both richness and turnover, the diversity of other, co-occurring taxa provides additional predictive power not captured by the environment alone. Our results suggest an important role for the biotic environment in future efforts aimed at modeling species assembly.

## Materials and Methods

Our goals were to assess the extent to which the taxonomic, phylogenetic, and functional dimensions of biodiversity and turnover reflect one another, and to parse the drivers of each dimension for plants, birds, and mammals across the USA. To do so, we used organismal data collected by NEON at 38 sites across the USA, climate data primarily reflecting temperature and water availability, published high-resolution phylogenies for each taxonomic group, and publicly accessible trait data on plant maximum height and mean bird and mammal body size. We estimated taxonomic, phylogenetic, and functional biodiversity within and turnover among sites, and then used these estimates to compare across diversity metrics within taxa, and examined the abiotic and biotic predictors of biodiversity and turnover for each taxonomic group. All analyses were conducted in R version 3.6.3 (R Core Team 2018) and all data and code are available on GitHub and also provided in the Supplement (Appendix S1).

### Organismal data

As part of the cross-site NEON protocol, scientists collect species occurrence data for plants, mammals, and birds at all sites (Thorpe et al. 2016). Although NEON site construction was officially completed in early 2019, site-level species occurrence data were collected at many NEON sites between 2012-2018. We used the ‘neonUtilities’ package (Lunch et al. 2020) to compile species inventories from three NEON Data Products: plant presence and percent cover (NEON.DP1.10058; NEON 2021), small mammal box trapping (NEON.DP1.10072; NEON 2021), and breeding landbird point counts (NEON.DP1.10003; NEON 2021). Of NEON’s 47 terrestrial field sites, 39 sites collected assemblage data for birds, plants, and mammals in 2017. We limited our data use to only those collected in 2017 to maximize overlap of assemblages in space and time, as NEON data offer a unique opportunity to compare local assemblages that connect at the regional scale. We removed one site (BARR: Utqiagvik, Alaska) from all analyses because only one mammal species was observed, resulting in 38 study sites distributed across the United States (Figure 1a). During the bootstrap sub-sampling of sites described below (see *Biodiversity Estimates*), estimates of mammal diversity were too low to estimate phylogenetic diversity in all but 13 sites; therefore, all analyses including mammal phylogenetic diversity or turnover as an explanatory or response variable use a reduced subset of sites (Appendix S2).

**Figure 1.**
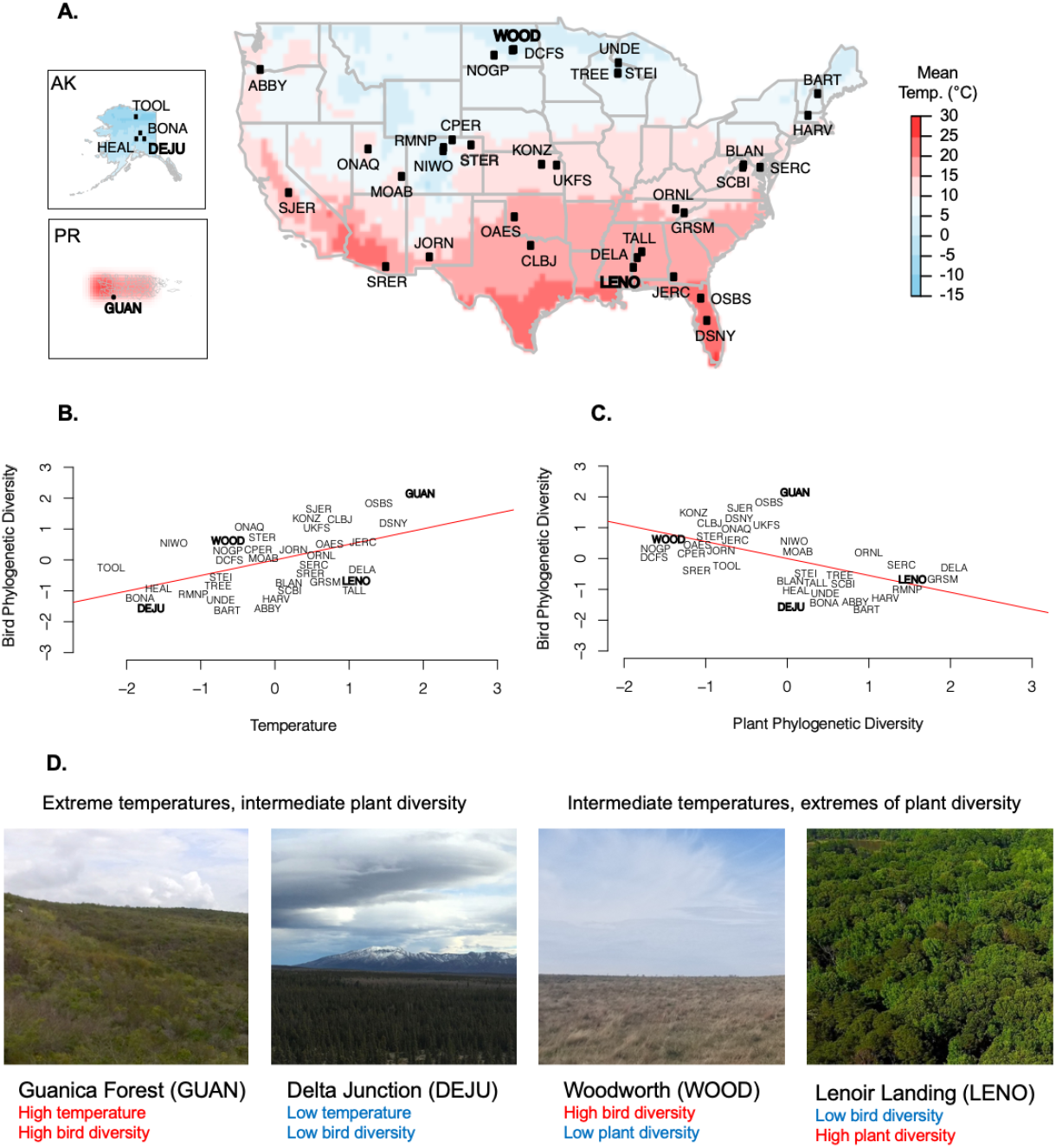
Abiotic and biotic predictors of bird phylogenetic diversity across NEON sites. (A) The locations of 38 NEON sites used in this study, at which plant, bird, and small mammal assemblages were surveyed in 2017. Colors represent mean annual temperatures (averaged across 30 years). Four sites, Guanica Forest (GUAN), Delta Junction (DEJU), Lenoir Landing (LENO) and Woodworth (WOOD) are highlighted to illustrate the relative contributions of predictors in subsequent panels. (B) Bird phylogenetic diversity has a positive relationship with temperature (or, the component axis capturing 75% of the variation in climate variables with strong loadings for temperature variables; red line is a simple linear model). Two sites, GUAN and DEJU are highlighted as sites with extreme temperatures that fit this relationship. (C) Bird phylogenetic diversity has a negative relationship with plant phylogenetic diversity. This relationship explains additional variation in bird phylogenetic diversity for some sites, including the highlighted sites WOOD and LENO. (D) Photographs representing the contrast of sites with intermediate plant diversity, where temperature is a stronger predictor of bird diversity, and sites with intermediate temperatures, where plant diversity is a stronger predictor of bird diversity.

### Site characteristics and climate

In order to generate potential abiotic predictors of biodiversity across NEON sites, we first gathered estimates of the average elevation, latitude, and longitude of each site, provided by NEON. To capture the climatic variables reflecting energy availability and potential abiotic constraints on regional species occurrences, we then accessed data for a suite of temperature and water availability variables (monthly mean, minimum, and maximum temperatures, precipitation, vapor pressure, cloud cover, frost day frequency, and potential evapotranspiration) from the Climatic Research Unit gridded product CRU TS v.4.03 (Harris et al. 2014). The CRU spatial resolution of 0.5° met our goal of highlighting broad differences among sites and minimizing the potential effects of small-scale processing error. Similarly, since we aimed to describe broad climatic differences among sites that might affect species occurrence, including establishment and persistence of individuals, we averaged all climatic variables across the years 1989-2018 and used these thirty-year average values in all subsequent analyses. We used Principal Component Analysis (PCA) to reduce the CRU climate variables to two principal components explaining a cumulative 93% of the variation in all abiotic predictors (Appendix S3). PC1 explained 75% of the total variation, with primary loadings from the three temperature variables, frost day frequency, vapor pressure, and potential evapotranspiration. The major loadings on PC2 include precipitation and cloud cover. We inverted the values of PC1 and refer to this axis as “temperature” and PC2 as “precipitation” for all subsequent analyses.

### Phylogeny

To estimate phylogenetic richness and turnover, we used published, high-resolution phylogenies for birds (Jetz et al. 2012b), mammals (Upham et al. 2019), and plants (Zanne et al. 2014). To account for phylogenetic uncertainty, we used a bootstrap approach for all phylogenetic analyses, estimating phylogenetic diversity measures and phylogenetic signal across 100 trees (following (Nakagawa and De Villemereuil 2019), who suggest that 50 would be sufficient) for each taxon. Bird and mammal trees were downloaded from VertLife.org, using species lists from the NEON organismal data (367 bird species, 93 mammal species). The bird trees were sampled from trees constrained by the backbone described in (Ericson et al. 2006) and contain 9993 OTUs each. The mammal trees were sampled from bird-death node-dated completed trees including 5911 species. To introduce plant species present in the organismal data but missing from the trees published in Zanne *et al*. (2014), we used the ‘congeneric.merge’ function in the package ‘pez’ (Pearse et al. 2015). This increased the species included in the plant phylogeny from 1667 to 2595 species.

### Trait Data

For estimates of functional richness and turnover, we used maximum plant height, average bird body mass, and average mammal body mass. Plant height data were gathered using the BIEN 4 trait database using the function ‘BIEN_trait_traitbyspecies’ in the package ‘BIEN’ (Maitner 2020), and used to calculate maximum plant height for 938 plant species. Mean bird body mass and mammal body mass values were gathered using the EltonTraits 1.0 database (Wilman et al. 2014) for 92 mammal species and 367 bird species. All traits were log transformed (natural log) to avoid bias in functional diversity from extreme trait values. For estimates of site-level functional diversity, we imputed trait values for the 1657 plant species and 1 mammal species for which we had no trait data, using the ‘phylopars’ function in the package ‘Rphylopars’ (Goolsby et al. 2017) and assuming a Brownian Motion model of trait evolution. This phylogenetic imputation is conservative, given our hypotheses are about the disconnects between functional and phylogenetic information, since it should tend to bias us towards finding similarities between functional and phylogenetic information. To estimate phylogenetic signal in each of the traits, we used the ‘phylosig’ function in the package ‘phytools’ v0.6.99 (Revell 2012), applied only to the non-imputed (i.e., truly measured) data.

### Biodiversity Estimates

For each taxonomic group (plants, birds, and mammals), we estimated measures of taxonomic, phylogenetic, and functional richness at, and turnover across, each of the 38 NEON sites, in order to identify the extent to which these metrics may reflect shared or unique processes.

While sampling protocols were kept consistent across sites, the number of plots sampled and number of visits to each plot differed. To standardize sampling across sites, within each taxon we used a bootstrap re-sampling approach to generate 100 estimates of each biodiversity metric. Each estimate was based on a random re-sampling of the minimum number of plots across sites (10 plots for plants, 5 for birds, 2 for mammals) and the minimum number of dates on which plots were sampled (1 date for all taxa). We calculated the mean of the 100 estimates for each biodiversity metric, resulting in one estimate for each taxon at each site. We compared these estimates to others calculated from all available data collected in 2017 for each site and found patterns of taxonomic, phylogenetic, and functional diversity to be comparable across sites (Appendix S4).

Taxonomic richness was calculated as total species richness, or the sum of unique species of each group (plants, mammals, birds), at each site. The phylogenetic richness at each site was estimated using the standardized effect size of mean nearest taxon distance, (SES_MNTD_; Webb et al. 2002, Kembel 2009) which reflects the extent to which species tend to be phylogenetically related to their nearest relative (and should be more sensitive to differences related to traits evolving under Brownian motion-type models; Letten and Cornwell 2015), while accounting for species richness. SES_MNTD_ was calculated using the function ‘ses.mntd’ in the package ‘picante’ v1.8.1 (Kembel et al. 2010). Functional richness was calculated as functional dispersion (Laliberté and Legendre 2010) using the function ‘dbFD’ in the ‘FD’ package (Laliberté et al. 2014) and was conducted using log-transformed maximum plant height and mean bird and mammal body mass.

Taxonomic turnover among sites is represented by Sörensen’s Index, which reflects the presence or absence of shared species between assemblages. We calculated Sörensen’s Index using the ‘vegdist’ function in the package ‘vegan’ v2.4-2 (Oksanen et al. 2019). We calculated turnover in phylogenetic diversity as PCDp (Ives and Helmus 2010), which isolates the average phylogenetic distance of non-shared species from the number of species shared in two communities (i.e., Sörensen’s index), using the ‘pez.dissimilarity’ function in the package ‘pez’ (Kembel et al. 2010, Pearse et al. 2015). We estimated turnover in functional diversity among sites using the same package, which uses a distance matrix of species’ trait values to estimate differences among sites.

### Statistical analysis

To compare taxonomic, phylogenetic, and functional diversity within each taxon, we used Pearson correlations for richness and Mantel tests for turnover (resulting in 9 analyses each for richness and turnover). In all models, data were Z-transformed to produce coefficients that reflect effect sizes (Gelman and Hill 2006).

To parse the abiotic and biotic predictors of taxonomic, phylogenetic, and functional richness, we used multiple linear regression models. For each diversity metric of each taxon (separately), we modeled diversity as a function of abiotic (temperature, precipitation, and elevation) and biotic explanatory variables (the same metric of diversity in the other two taxonomic groups; *i.e*. bird phylogenetic diversity as a function of plant and mammal phylogenetic diversity). To decompose variance explained by abiotic and biotic factors, we also compared two additional sets of models with only abiotic, and only biotic explanatory variables. We compared the R^2^ values of each model to assess the role of biotic and abiotic predictors across taxa and metrics of biodiversity and used the coefficients of the predictors from the full models to compare their strength and significance. Additional analyses to ensure correlations between abiotic and biotic factors did not affect our results are shown in the supplementary materials.

To estimate the relative impact of abiotic and biotic factors on taxonomic, phylogenetic, and functional turnover across NEON sites for plants, mammals, and birds, we used quantile regression models. These models are ideal for comparing multiple distance matrices, and while we include spatial distance in these analyses to control for autocorrelation we emphasize that quantile regressions are robust to even pseudoreplication (Legendre and Legendre 2012). As with the models of richness, for each diversity metric in each taxon we modeled diversity as function of abiotic and biotic explanatory variables, abiotic variables only, biotic variables only, and we modeled the residuals of the abiotic models as a function of the biotic predictors. The variables included were the same as those from the alpha-diversity models but were calculated as dissimilarity values between sites rather than site averages, and with distance among sites also included among the abiotic variables. We used the coefficients of the predictors from the full models to identify primary correlates of beta-diversity.

## Results

### Relationships among taxonomic, phylogenetic, and functional richness and turnover

Taxonomic, phylogenetic, and functional diversity and turnover were generally positively, but weakly, correlated with one another (Table 1). The strength and signal of associations among the metrics varied across taxa. We found the strongest associations among the phylogenetic and functional richness and turnover of plants and birds (richness; plants: r_36_ = 0.78, p < 0.001; birds: r_36_ = 0.49, p = 0.002; turnover; plants: r_701_ = 0.48, p < 0.001; birds: r _701_ = 0.32, p < 0.001).

**Table 1.**
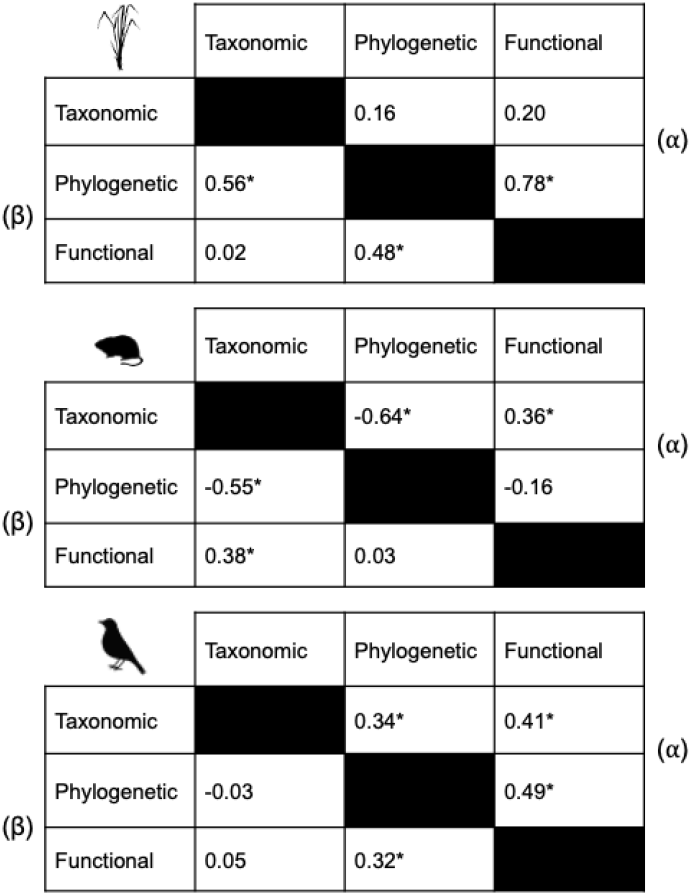
Correlation among taxonomic, phylogenetic, and functional richness and turnover for plants, mammals, and birds across NEON sites. We compared each biodiversity metric to all others to estimate the extent to which they captured similar biodiversity pattern and process across 38 sites in the US. Relationships among metrics differed between richness (upper right of each table, noted with α) and turnover (lower left of each table, noted with β) and among taxa, with stronger relationships between phylogenetic and functional richness and turnover for both plants and birds, as well as significant positive correlations between taxonomic and functional richness for birds and mammals, that held true for mammal turnover but not for birds.

|  |  |  |  |  |  |
| --- | --- | --- | --- | --- | --- |
|  |  | Taxonomic | Phylogenetic | Functional | (α) |
| (β) | Taxonomic |  | 0.16 | 0.20 |  |
|  | Phylogenetic | 0.56* |  | 0.78* |  |
|  | Functional | 0.02 | 0.48* |  |  |
|  |  | Taxonomic | Phylogenetic | Functional | (α) |
| (β) | Taxonomic |  | -0.64* | 0.36* |  |
|  | Phylogenetic | -0.55* |  | -0.16 |  |
|  | Functional | 0.38* | 0.03 |  |  |
|  |  | Taxonomic | Phylogenetic | Functional | (α) |
| (β) | Taxonomic |  | 0.34* | 0.41* |  |
|  | Phylogenetic | -0.03 |  | 0.49* |  |
|  | Functional | 0.05 | 0.32* |  |  |

Taxonomic and functional richness were positively correlated for mammals (r_36_ = 0.36, p = 0.03) and birds (r_36_ = 0.41, p = 0.01), as were taxonomic and functional turnover for mammals (r_701_ = 0.38, p < 0.001), but not birds (r_701_ = 0.05, p = 0.22). Lastly, taxonomic and phylogenetic richness, but not turnover, were positively correlated in birds (r_36_ = 0.34, p = 0.04), while these metrics were positively correlated for turnover, but not richness, in plants (r_701_ = 0.56, p < 0.001). For both richness and turnover, the same metrics were significantly, negatively correlated for mammals (richness; r_11_ = -0.64, p = 0.02; turnover; r_497_ = -0.55, p < 0.001).

### Predictors of plant, mammal and bird richness

Across all taxonomic groups and measures of biodiversity, the strongest models of richness included both abiotic and biotic explanatory variables (Figure 2). Taken together, abiotic and biotic factors explained 92%, 68%, and 85% of the variation in plant, mammal, and bird phylogenetic richness (respectively) across NEON sites. The majority of the variation in phylogenetic richness of plants and birds was explained by the biotic environment; this was also the case in model fits that first accounted for abiotic variables (Appendix S5).

**Figure 2.**
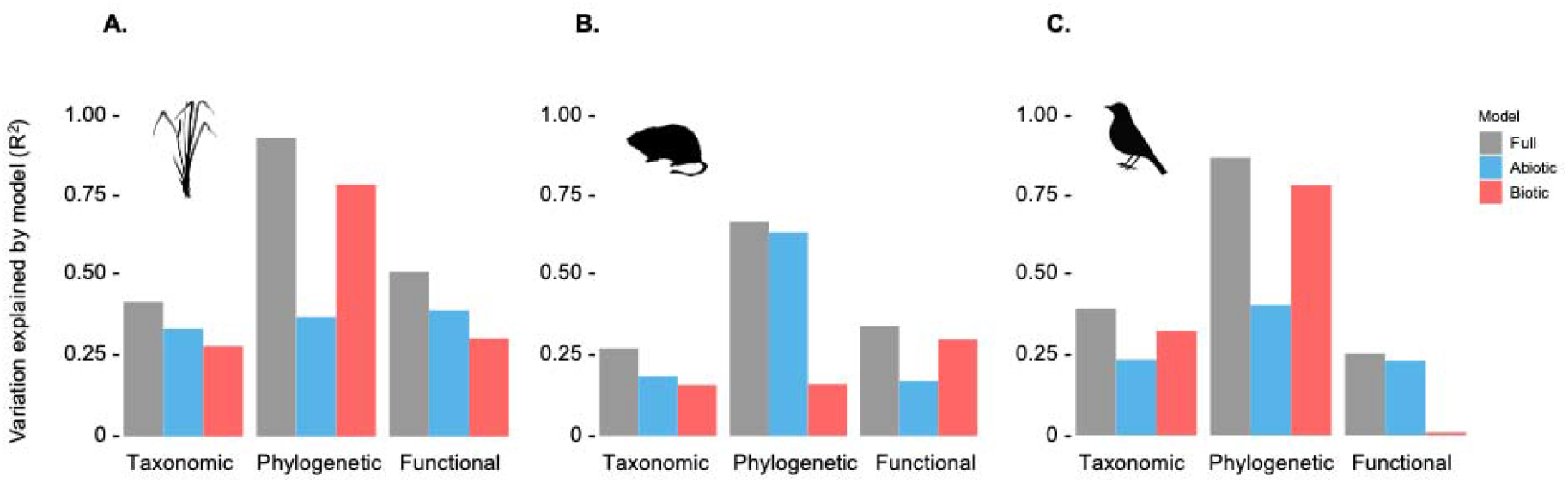
Explanatory power of abiotic, biotic, and combined models of biodiversity richness. We report R-squared values from multiple linear regression models, including ‘full’ models using all abiotic predictors (PC1, PC2, elevation) and biotic predictors (corresponding biodiversity measures of other taxa), models using only abiotic predictors, and models using only biotic predictors. We present results for models of taxonomic, phylogenetic, and functional diversity of (A) plants, (B) mammals, and (C) birds.

Among the abiotic predictors considered here, we found that either temperature, precipitation, or both tended to have positive associations with plant, bird, and mammal richness (Figure 3a). For plants, temperature was a significant driver of species richness (p = 0.04, df = 32) and had a strong positive association with phylogenetic richness (p = 0.08, df = 7), while precipitation was a significant driver of both phylogenetic (p = 0.05, df = 7) and functional richness (p = 0.02, df = 32). Similarly, although non-significant, bird phylogenetic richness tended to be higher at warmer (Figure 1b; p = 0.15, df = 7), wetter sites (p = 0.35), and mammal phylogenetic diversity was higher at wetter sites (p = 0.21, df = 7). Bird and mammal functional diversity were more poorly predicted by the abiotic environment, however bird functional diversity had a significant negative relationship with elevation; that is, higher elevation sites tended to have lower bird functional richness (p = 0.04, df = 32).

**Figure 3.**
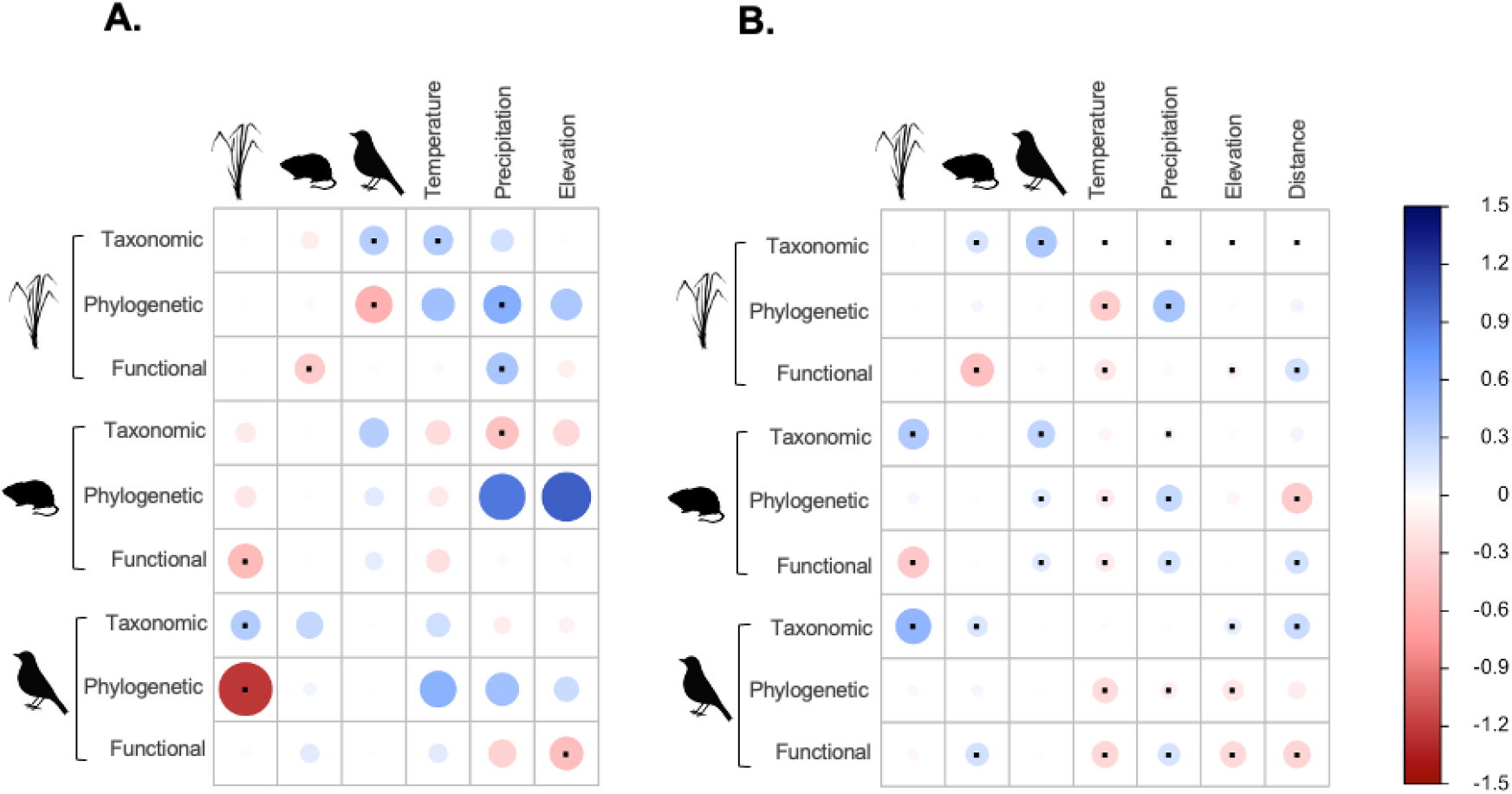
The abiotic and abiotic predictors of plant, mammal, and bird richness and turnover. Panels reflect coefficients from models of (A) richness (linear regression), and (B) turnover (quantile regression), where central points indicate significance (p < 0.05). Each row reflects one model and each column one predictor; biotic predictors reflect the same biodiversity metric as the response variable (*e.g*., models of mammal phylogenetic diversity have inputs of bird and plant phylogenetic diversity). (A) models of biodiversity richness show plant diversity is strongly associated with temperature, and the phylogenetic and functional diversity of birds and mammals are negatively associated with that of plants. (B) models of biodiversity turnover show strong cross-taxon biotic associations even after accounting for the abiotic environment. All data were scaled, resulting in coefficients that reflect relative (standard) effect sizes.

Among the biotic predictors of species richness, we found a significant, positive association between plant and bird taxonomic diversity (p = 0.04, df = 32). We also found a significant, negative association between the phylogenetic diversity of plants and birds (p = 0.04, df = 7) as well as between the functional diversity of plants and mammals (p = 0.01, df = 32). That is, although sites with more plant species also tend to host more bird species, those with higher phylogenetic and functional plant richness also tended to host more clustered communities of birds and mammals, and vice versa (Figure 1c-d).

### Predictors of plant, mammal and bird turnover

The predictors of turnover in species identities, clades, and functions also included contributions from both the abiotic and biotic environment (Figure 3b). In many cases the strongest predictors of turnover reflected turnover in other co-occurring taxa. For example, turnover in the species of birds and mammals (taxonomic turnover) among sites was best predicted by that of plants (both p < 0.001, df = 703) even when accounting for abiotic differences and distances among sites (Figure 4, Appendix S6). However, this was not the case for phylogenetic turnover in either consumer taxon (birds: p = 49, df = 499; mammals: p = 0.21, df = 499), or for functional turnover in birds (p = 0.14, df = 703). Rather, differences in precipitation among sites showed positive associations with phylogenetic turnover in plants and mammals (p < 0.001, df = 499) and functional turnover in mammals and birds (p < 0.001, df = 703). That is, sites that differed the most in precipitation tended to host assemblages that were more phylogenetic and functionally distinct from one another. The opposite was true of temperature; with greater differences in temperature among sites, assemblages tended to have greater similarities in the clades and functions represented (Figure 3b).

**Figure 4.**
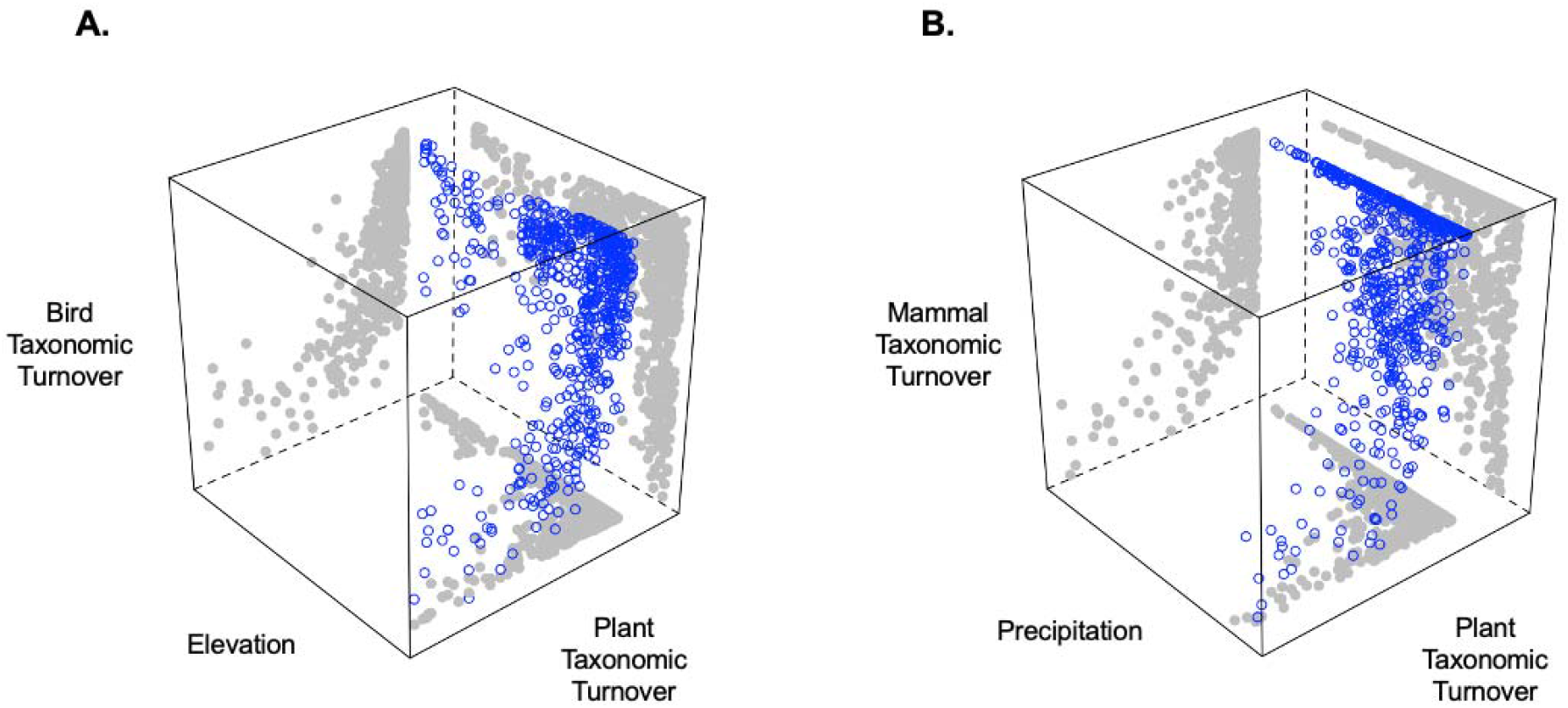
Consumer species turnover is best predicted by plant turnover, followed by environmental differences among sites. (A) Bird species turnover is strongly predicted by plant turnover, and more weakly predicted by differences in elevation among sites. (B) Mammal species turnover is best predicted by differences in plant turnover, followed by differences in precipitation among sites.

## Discussion

Here we applied plant and animal assemblage data collected by NEON to investigate the patterns and drivers of taxonomic, phylogenetic, and functional richness and turnover across the USA. NEON’s organismal data uniquely captures records of co-occurring plants, mammals, and birds at a fine spatial grain (reflecting local-scale ecological dynamics) spread across a broad spatial extent (at which broad-scale evolutionary processes take place). Across NEON sites, we compared these taxonomic, phylogenetic, and functional richness and turnover within taxa and found that they were often weak predictors of one another. We also parsed the abiotic and biotic drivers of richness and turnover for each taxon, and found support for the role of the biotic environment in shaping species assembly, even when accounting for variation in the abiotic environment (*e.g*., energy inputs).

### General patterns and correlates among biodiversity facets

The often-weak relationships we detected among taxonomic, phylogenetic, and functional diversity and turnover reflect unique patterns and processes. Among the strongest associations of these metrics were between phylogenetic and functional richness in plants and birds. This is expected to occur when evolutionary distance between species is representative of trait and niche differentiation (Webb et al. 2002, Cadotte et al. 2009, Devictor et al. 2010), and so traits representing function have strong phylogenetic signal, as is true for all traits included in this study (all Pagel’s lambda > 0.95, all p < 0.001; Pagel 1999). The stronger correlation between phylogenetic and functional diversity in plants reflects widespread phylogenetic signal in plant traits (Pennell et al. 2015), while limitations on the strength of this relationship in the consumer taxa echo previous findings of inconsistency of species richness as a predictor of phylogenetic diversity over space and time (Devictor et al. 2010, Flynn et al. 2011, Purschke et al. 2013). In particular, while previous studies have similarly found positive associations between phylogenetic and functional diversity for birds (Pigot et al. 2016), the convergent evolution of morphological traits (Pigot et al. 2020) and the reversal of global trait patterns within lineages (Nee et al. 1991) can limit the ability of phylogenetic diversity metrics to inform function.

We found a positive relationship between taxonomic and functional diversity in mammals and birds, suggesting that species-rich consumer assemblages typically reflect a broad range of function. Considering the traits analyzed all exhibit phylogenetic signal, it is somewhat surprising that taxonomic and phylogenetic richness (measured as SES_MNTD_, which controls for null effects of species richness) are significantly positively associated only in birds. This difference among taxa may reflect lesser dispersal in plants: if mammals and birds assemble from a broader regional pool, and thus from more phylogenetic lineages, then the addition of species may be more likely to result in increased evolutionary variety, as compared to dispersal-limited producers. Links between bird taxonomic and phylogenetic alpha-diversity have been demonstrated using data with a broad grain and extent (Voskamp et al. 2017), while higher resolution data sets have shown weak relationships between the two metrics (Devictor et al. 2010), as is the case here. This demonstrates the importance of using fine-grain data for exploring the processes underlying site-level biodiversity and suggests these measures will contribute somewhat independently to our understanding of alpha-diversity.

Lastly, taxonomic diversity had a significant, positive relationship with PCDp for plants, but not for birds or mammals; as the proportion of plant species that are shared among assemblages increases, so does the phylogenetic distance between non-shared species. We might expect this to be the case if, in two communities with few species in common, close relatives occupy similar niches (in which case species not in common would be closely related). The significant negative relationship in mammals indicates that as communities have more species in common, non-shared species are more closely related. This may reflect a more regional-scale pattern, wherein communities that include many of the same species (*e.g*., because they share a biome) will also contain closely-related, non-shared species due to shared biogeography (Graham and Fine 2008).

### Abiotic and biotic environment together predict richness

The strength of individual aspects of the biotic and abiotic environment at predicting plant, bird, and mammal diversity suggest joint contributions of broad- and local-scale processes. While abiotic predictors reflecting broad-scale patterns in energy inputs consistently explained some variation in biodiversity, as in previous studies (Currie et al. 2004, Jetz et al. 2008, Meynard et al. 2011), in many cases the biotic environment alone explained as much or more variation. This suggests that at the broad scale, energy inputs influence processes like species radiation, while at the local-scale (particularly at sites with intermediate energy inputs) local-scale processes of niche diversification explain additional variation (Graham et al. 2018). Bird and plant taxonomic diversity were positively associated with one another, consistent with previously published global and regional patterns (Gaston 2000, Kissling et al. 2007, 2008). It is widely recognized that changes in the diversity of one taxonomic group can result in similar changes to another by altering available niche space through resources and interactions (Kissling et al. 2007, 2008, Barrio et al. 2016). However, our findings contrast with some studies of biodiversity over broad spatial extents (and grains) which have concluded that the variation in biodiversity explained by other taxa is largely redundant to that explained by the abiotic environment (Jetz et al. 2008).

One example of the power of the biotic environment is our finding that phylogenetic bird and plant richness were better predictors of one-another than was the abiotic environment. A site with few closely-related plant species (*e.g*., a grassland) was likely to host a larger phylogenetic diversity (corrected for species richness) of birds (and functional diversity of mammals), while a site with more distantly-related plant species (*e.g*., a highly-structured forest) was more likely to contain species representing young, rapidly diversifying clades (*e.g*., passerines) and therefore a lower phylogenetic diversity of birds. Bird functional diversity, on the other hand, was best predicted by the abiotic predictor of elevation, consistent with past studies that have shown strong elevational effects on diversification (McCain 2009, Quintero and Jetz 2018). Declines in taxonomic diversity with elevation, as seen here, can result from shifts in plant structure and ecosystem productivity (Kattan and Franco 2004, Herzog et al. 2005).

### The biotic environment is a valuable predictor of turnover

Congruence in the turnover of different taxa has previously been identified for plants, invertebrates, mammals, and birds over broad cross-continental spatial extents (McKnight et al. 2007, Buckley and Jetz 2008), and local to regional extents (Su et al. 2004, Oertli et al. 2005, Steinitz et al. 2005); however, whether turnover is driven primarily by turnover in other taxa (*e.g*., Buckley and Jetz 2008) or climate (*e.g*., Zellweger et al. 2017) has been unclear. Our results, which are unique in that they reflect assemblages of species which co-occur at the local scale, across taxonomic groups, located across regional environmental gradients, suggest the best models of plant and mammal turnover incorporate both abiotic and biotic predictors.

The strength of the cross-taxon biotic environment for predicting turnover (even after accounting for broad-scale environmental variation) highlights the potential of site-level biotic processes to drive species turnover. Our results specifically reflect that taxonomic turnover in producer assemblages predicts turnover in consumers. Links between the composition of plants and birds are particularly well-demonstrated (Lee and Rotenberry 2005, Fleishman and Mac Nally 2006, Jankowski et al. 2013) and while foraging relationships are often implicated, vegetation structure can also be an important mediator between plant and bird turnover (Zellweger et al. 2017). Our results regarding functional turnover suggest that if this is the case, maximum plant height may capture only a small portion of the features of vegetative structure most relevant to variation in bird body size.

The importance of the biotic environment to turnover compared to local diversity reflects the processes driving the composition (turnover), as compared to the local richness, of biodiversity. The addition or loss of particular producer species can lead to corresponding changes in consumers that rely on those species for resources and services; the same can be true for clades of related species when close relatives share similar roles (Graham and Fine 2008).

## Conclusions

Biodiversity conservation interventions are often managed and assessed at the site-level, but site-level diversity results from broad-scale dynamics like speciation and biogeography, in addition to local-scale dynamics like environmental filtering and biotic interactions. Studies capturing those broad-scale dynamics tend to be limited to broad spatial grains (*e.g*., using species distributions to estimate co-occurrence), while studies that do capture the fine grain at which local-scale dynamics operate tend to be limited in their extent. Using fine grain species inventories for plants, mammals, and birds from NEON spread out over the broad extent of the United States, enabled us to investigate the drivers of site-level biodiversity in a new light.

We found that both the abiotic and biotic environments explained separate aspects of biodiversity and turnover. We found nuanced relationships in taxonomic and phylogenetic diversity, particularly for plants and birds, which reflect a combination of broad-scale species processes of speciation along abiotic gradients, and local-scale niche diversification leading to an inverse relationship between the taxa. Most strikingly, we found that plant turnover among sites is a stronger predictor of bird and mammal taxonomic turnover than individual environmental differences between sites, suggesting shifts in producer diversity can predict consistent changes in consumer turnover across sites, habitats, and biomes that are not explained by shared abiotic responses.

While associations among taxa may reflect shared environmental responses not captured by the abiotic or biotic variables included here (*e.g*., land use and microhabitat features, invertebrate biodiversity), it is also widely recognized that taxonomic groups are not isolated from one another in processes of species assembly, and through niche dynamics biodiversity can beget biodiversity (Burghardt et al. 2009). Only through continued investigation of the role of the biotic environment, by parsing the ecological and evolutionary patterns among species’ environmental tolerances and co-occurrences within and across taxa (*e.g*., with tools like phylogenetic generalized linear mixed models; (Gallinat and Pearse 2021), can we better understand the role of the abiotic and biotic environments in fundamental ecology as well as conservation practice.

## Acknowledgments

We thank NEON staff, particularly Kate Thibault and Clare Lunch, for their help with the use of NEON data. ASG, WDP, and the Pearse Lab are funded by National Science Foundation grants ABI-1759965 and EF-1802605, and UKRI/NERC NE/V009710/1. The National Ecological Observatory Network is a program sponsored by the National Science Foundation and operated under cooperative agreement by Battelle. This material is based in part upon work supported by the National Science Foundation through the NEON Program.

## Appendices

**Appendix S1**. All R scripts and data products used for analysis are available at https://github.com/gallinamanda/neon-biodiversity and will be included in the supplement as a .zip file.

**Appendix S2**. Site information, including site code, name, state, latitude (lat), and longitude (lon), with relative (scaled) measures of taxonomic (tax), phylogenetic (phy), and functional (fun) richness for plants, mammals, and birds. Estimates are based on a bootstrap subsampling approach (see *Biodiversity Estimates* in Materials and Methods) to standardize sampling among sites. For 25 of 38 sites, this approach resulted in too few mammal species to calculate phylogenetic diversity (those without estimates are labeled “NA” below).

**Appendix S3a**. Summary of importance of components in PCA describing the abiotic environmental descriptors of 38 NEON sites. Variables included monthly mean, minimum, and maximum temperatures, precipitation, vapor pressure, cloud cover, frost day frequency, and potential evapotranspiration, accessed from the Climatic Research Unit gridded product CRU TS v.4.03 (Harris et al. 2014). All climatic variables were averaged across the years 1989-2018.

**Appendix S3b**. Loadings of abiotic environmental variables onto two principal components.

**Appendix S4**. Comparison of estimates of biodiversity from re-sampled data and all available data. Species inventories for plants, mammals, and birds were sampled unevenly across 38 NEON sites in 2017; the number of plots surveyed and frequency of sampling at each plot differed across and within sites. We used a bootstrap sampling approach to re-sample assemblages 100 times based on the minimum number of plots sampled at each site (plants: 10, birds: 5, mammals: 2) and minimum number of dates during which each plot was sampled (1 for all taxa). Here we compare the average taxonomic, phylogenetic, and functional diversity values for each taxon, averaged from 100 bootstrap values, with the same diversity measures calculated using all available species occurrence data regardless of plot and date visitation. Red lines indicate a 1:1 relationship. Taxonomic, phylogenetic, and functional diversity all follow similar patterns in both data sets, with taxonomic diversity being lower in the bootstrap values, as is expected. Phylogenetic and functional diversity control for the number of species, resulting in a relationship closer to 1:1 between the two data sets. In all cases, relationships between the bootstrap values and all-data values are weakest for mammals. This is likely due to there being fewer mammal species overall compared to plants and birds, causing species absences during re-sampling to result in large differences in phylogenetic and functional composition.

**Appendix S5**. Explanatory power (R-squared values, reported here as percentages) of richness models, including models of abiotic residuals predicted by biotic predictors. Values describe linear regression models including full models using all abiotic predictors (PC1, PC2, elevation), and biotic predictors (corresponding biodiversity measures of other taxa), models using only abiotic predictors, models using only biotic predictors, and abiotic model residuals as a function of biotic predictors (labeled “residual” in the table). We present results for models of taxonomic, phylogenetic, and functional diversity of plants, mammals, and birds.

**Appendix S6**. Residuals from mammal and bird abiotic turnover models as a function of plant turnover. For each diversity metric, including taxonomic diversity (Sorensen’s Index, left), phylogenetic diversity (PCDp, middle panels), and functional diversity (Functional Dissimilarity, right), residuals from models using only abiotic explanatory variables (environmental PC1, PC2, and elevation) are plotted as a function of plant diversity.

