## Appendices S1-S6 for "The abiotic and biotic environment together predict plant, mammal, and bird diversity and turnover across the United States"

Appendix S1. All R scripts and data products used for analysis are available at <https://github.com/gallinamanda/neon-biodiversity> and will be included in the supplement as a .zip file.

Appendix S2. Site information, including site code, name, state, latitude (lat), and longitude (lon), with relative (scaled) measures of taxonomic (tax), phylogenetic (phy), and functional (fun) richness for plants, mammals, and birds. Estimates are based on a bootstrap subsampling approach (see *Biodiversity Estimates* in Materials and Methods) to standardize sampling among sites. For 25 of 38 sites, this approach resulted in too few mammal species to calculate phylogenetic diversity (those without estimates are labeled “NA” below).

| Site | Name | State | Lat | Lon | Plant  Tax | Plant  Phy | Plant  Fun | Bird  Tax | Bird  Phy | Bird  Fun | Mammal  Tax | Mammal  Phy | Mammal  Fun |
| --- | --- | --- | --- | --- | --- | --- | --- | --- | --- | --- | --- | --- | --- |
| ABBY | Abby Road | WA | 45.76 | -122.33 | -0.17 | 0.84 | -0.32 | -0.60 | -1.38 | -1.05 | 1.47 | 0.52 | 0.82 |
| BART | Bartlett Experimental Forest | NH | 44.06 | -71.29 | -1.66 | 0.98 | 2.73 | -0.54 | -1.61 | -1.81 | 0.80 | 1.57 | -0.57 |
| BLAN | Blandy Experimental Farm | VA | 39.06 | -78.07 | 1.01 | 0.05 | 0.25 | -0.55 | -0.85 | -1.37 | -0.34 | NA | -0.88 |
| BONA | Caribou-Poker Creeks Research Watershed | AK | 65.15 | -147.50 | -1.28 | 0.45 | 0.88 | -1.17 | -1.22 | -0.37 | -0.76 | NA | 0.32 |
| CLBJ | Lyndon B. Johnson National Grassland | TX | 33.40 | -97.57 | 1.85 | -0.95 | -0.35 | 0.90 | 1.17 | 1.20 | 1.29 | -1.25 | 1.73 |
| CPER | Central Plains Experimental Range | CO | 40.82 | -104.75 | -0.55 | -1.17 | -1.27 | -1.26 | 0.34 | -1.57 | -0.81 | NA | 1.32 |
| DCFS | Dakota Coteau Field | ND | 47.16 | -99.11 | -0.14 | -1.63 | -1.38 | -0.74 | 0.19 | 1.02 | -0.71 | NA | -0.45 |
| DEJU | Delta Junction | AK | 63.88 | -145.75 | -0.82 | 0.05 | 0.14 | -1.00 | -1.48 | -0.05 | -1.06 | NA | 0.46 |
| DELA | Dead Lake | AL | 32.54 | -87.80 | 0.85 | 2.05 | 1.01 | -0.66 | -0.29 | 0.71 | -1.17 | NA | -1.36 |
| DSNY | Disney Wilderness Preserve | FL | 28.13 | -81.44 | -0.14 | -0.59 | 0.04 | 0.22 | 1.20 | 0.95 | -0.82 | NA | 0.30 |
| GRSM | Great Smoky Mountains National Park | TN | 35.69 | -83.50 | 0.66 | 1.91 | 1.55 | 0.49 | -0.65 | -0.33 | -0.73 | NA | -2.01 |
| GUAN | Guanica Forest | PR | 17.97 | -66.87 | -0.08 | 0.09 | -0.57 | 0.23 | 2.15 | 0.79 | -1.21 | NA | 0.68 |
| HARV | Harvard Forest & Quabbin Watershed | MA | 42.54 | -72.17 | -0.34 | 1.21 | 1.35 | 0.94 | -1.14 | -1.39 | 0.80 | -0.04 | -0.01 |
| HEAL | Healy | AK | 63.88 | -149.21 | -1.25 | 0.22 | 0.19 | -1.17 | -0.90 | 0.23 | -0.49 | NA | 0.41 |
| JERC | The Jones Center at Ichauway | GA | 31.19 | -84.47 | 1.36 | -0.70 | -0.07 | 0.45 | 0.60 | -0.22 | -0.19 | NA | 0.21 |
| JORN | Jornada Experimental Range | NM | 32.59 | -106.84 | -1.11 | -0.84 | -0.72 | -0.62 | 0.36 | 0.60 | 1.66 | -0.24 | 1.61 |
| KONZ | Konza Prairie Agroecosystem | KS | 39.10 | -96.56 | 1.41 | -1.15 | -0.53 | 1.24 | 1.36 | 0.33 | 1.54 | 0.18 | 1.11 |
| LENO | Lenoir Landing | AL | 31.85 | -88.16 | 0.47 | 1.53 | 1.04 | -0.78 | -0.76 | 0.77 | -1.23 | NA | -1.61 |
| MOAB | Moab | UT | 38.25 | -109.39 | -1.50 | 0.13 | -0.67 | -0.13 | 0.23 | -0.45 | 0.15 | NA | 0.00 |
| NIWO | Niwot Ridge | CO | 40.05 | -105.58 | -0.12 | 0.09 | -0.48 | -0.47 | 0.57 | -1.02 | -0.55 | NA | 0.83 |
| NOGP | Northern Great Plans Research Laboratory | ND | 46.77 | -100.92 | -0.34 | -1.61 | -1.30 | -0.55 | 0.33 | -0.15 | -0.53 | NA | 0.78 |
| OAES | Marvin Klemme Range Research Station | OK | 35.41 | -99.06 | 0.28 | -1.10 | -0.96 | 0.85 | 0.46 | 0.53 | 0.57 | -0.70 | 0.49 |
| ONAQ | Onaqui | UT | 40.18 | -112.45 | -1.44 | -0.62 | -1.62 | 0.03 | 1.07 | 0.05 | 0.63 | NA | -0.08 |
| ORNL | Oak Ridge | TN | 35.96 | -84.28 | 1.29 | 1.00 | 1.27 | 1.67 | 0.20 | 0.79 | -0.78 | NA | -0.92 |
| OSBS | Ordway-Swisher Biological Station | FL | 29.69 | -81.99 | 0.04 | -0.22 | 0.43 | 0.02 | 1.85 | 2.17 | -0.68 | NA | -0.29 |
| RMNP | Rocky Mountain National Park | CO | 40.28 | -105.55 | 0.09 | 1.47 | -0.19 | -0.34 | -0.96 | -0.54 | -0.95 | NA | -0.24 |
| SCBI | Smithsonian Conservation Biology Institute | VA | 38.89 | -78.14 | 1.60 | 0.67 | 0.13 | -0.45 | -0.99 | -1.71 | -0.25 | NA | -1.47 |
| SERC | Smithsonian Environmental Research Center | MD | 38.89 | -76.56 | 0.59 | 1.40 | 1.06 | 0.05 | -0.19 | -0.13 | -1.03 | NA | -1.78 |
| SJER | San Joaquin Experimental Range | CA | 37.11 | -119.73 | -0.43 | -0.58 | -0.67 | 2.11 | 1.65 | 1.14 | 0.09 | NA | -0.96 |
| SRER | Santa Rita Experimental Range | AZ | 31.91 | -110.84 | -0.45 | -1.11 | -0.43 | 0.29 | -0.37 | 0.75 | 2.77 | -1.87 | 0.66 |
| STEI | Steigerwaldt-Chequamegon | WI | 45.51 | -89.59 | -0.12 | 0.23 | 0.97 | 1.13 | -0.59 | -0.06 | 0.95 | -0.67 | -1.19 |
| STER | North Sterling | CO | 40.46 | -103.03 | -1.61 | -0.94 | -1.75 | -1.73 | 0.74 | -2.14 | 0.13 | 0.34 | -0.18 |
| TALL | Talladega National Forest | AL | 32.95 | -87.39 | 1.92 | 0.35 | 0.75 | 0.87 | -0.77 | -0.47 | -0.69 | NA | -1.18 |
| TOOL | Toolik Field Station | AK | 68.66 | -149.37 | -0.65 | -0.74 | -0.63 | -2.05 | -0.22 | -0.22 | -0.59 | 1.73 | 1.15 |
| TREE | Treehaven | WI | 45.49 | -89.59 | 0.03 | 0.49 | 0.66 | -0.91 | -0.79 | 0.42 | 1.04 | 0.17 | -0.43 |
| UKFS | KU Field Station | KS | 39.04 | -95.19 | 1.57 | -0.28 | -0.24 | 1.63 | 1.10 | 0.29 | 0.00 | NA | 1.61 |
| UNDE | University of Notre Dame Environmental Research Center | MI | 46.23 | -89.54 | -0.32 | 0.47 | 0.95 | 1.36 | -1.04 | 0.53 | 1.74 | 0.25 | 0.23 |
| WOOD | Chase Lake National Wildlife Refuge | ND | 47.13 | -99.24 | -0.49 | -1.45 | -1.25 | 1.22 | 0.63 | 1.77 | -0.06 | NA | 0.88 |

Appendix S3a. Summary of importance of components in PCA describing the abiotic environmental descriptors of 38 NEON sites. Variables included monthly mean, minimum, and maximum temperatures, precipitation, vapor pressure, cloud cover, frost day frequency, and potential evapotranspiration, accessed from the Climatic Research Unit gridded product CRU TS v.4.03 (Harris et al. 2014). All climatic variables were averaged across the years 1989-2018.

|  | PC1 | PC2 | PC3 | PC4 | PC5 | PC6 | PC7 | PC8 |
| --- | --- | --- | --- | --- | --- | --- | --- | --- |
| standard deviation | 2.44 | 1.22 | 0.58 | 0.33 | 0.28 | 0.14 | 0.07 | <0.01 |
| Proportion of variance | 0.75 | 0.19 | 0.04 | 0.01 | 0.01 | <0.01 | <0.01 | <0.01 |
| Cumulative proportion | 0.75 | 0.93 | 0.97 | 0.99 | 1.00 | 1.00 | 1.00 | 1.00 |

Appendix S3b. Loadings of abiotic environmental variables onto two principal components.

| Abiotic variable | PC1 | PC2 |
| --- | --- | --- |
| Mean temperature | -0.41 | -0.05 |
| Minimum temperature | -0.41 | 0.06 |
| Maximum temperature | -0.40 | -0.15 |
| Precipitation | -0.30 | 0.51 |
| Vapor pressure | -0.37 | 0.30 |
| Potential evapotranspiration | -0.33 | -0.31 |
| Cloud cover | 0.16 | 0.73 |
| Frost day frequency | 0.38 | -0.03 |

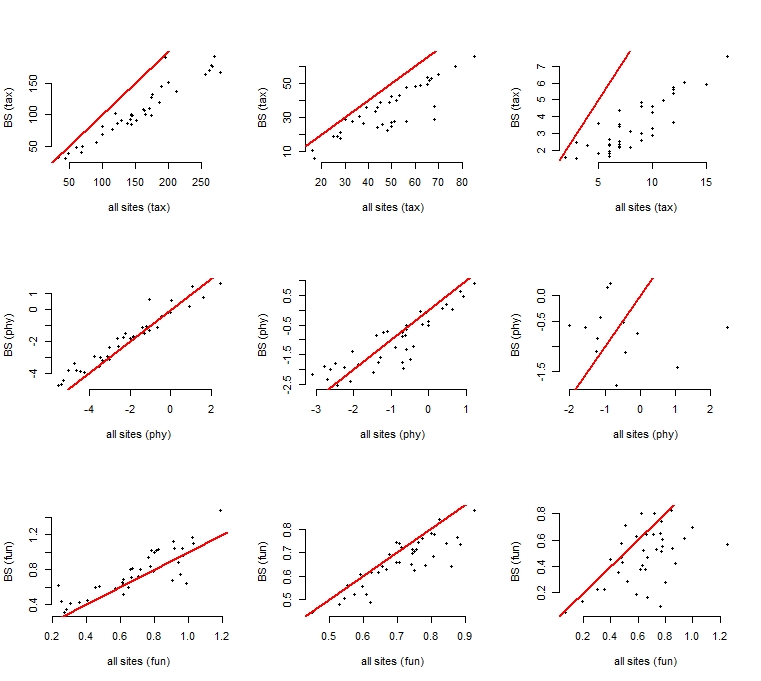

Appendix S4. Comparison of estimates of biodiversity from re-sampled data and all available data. Species inventories for plants, mammals, and birds were sampled unevenly across 38 NEON sites in 2017; the number of plots surveyed and frequency of sampling at each plot differed across and within sites. We used a bootstrap sampling approach to re-sample assemblages 100 times based on the minimum number of plots sampled at each site (plants: 10, birds: 5, mammals: 2) and minimum number of dates during which each plot was sampled (1 for all taxa). Here we compare the average taxonomic, phylogenetic, and functional diversity values for each taxon, averaged from 100 bootstrap values, with the same diversity measures calculated using all available species occurrence data regardless of plot and date visitation. Red lines indicate a 1:1 relationship. Taxonomic, phylogenetic, and functional diversity all follow similar patterns in both data sets, with taxonomic diversity being lower in the bootstrap values, as is expected. Phylogenetic and functional diversity control for the number of species, resulting in a relationship closer to 1:1 between the two data sets. In all cases, relationships between the bootstrap values and all-data values are weakest for mammals. This is likely due to there being fewer mammal species overall compared to plants and birds, causing species absences during re-sampling to result in large differences in phylogenetic and functional composition.

Appendix S5. Explanatory power (R-squared values, reported here as percentages) of richness models, including models of abiotic residuals predicted by biotic predictors. Values describe linear regression models including full models using all abiotic predictors (PC1, PC2, elevation), and biotic predictors (corresponding biodiversity measures of other taxa), models using only abiotic predictors, models using only biotic predictors, and abiotic model residuals as a function of biotic predictors (labeled “residual” in the table). We present results for models of taxonomic, phylogenetic, and functional diversity of plants, mammals, and birds.

|  |  | Plants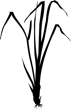 | Mammals 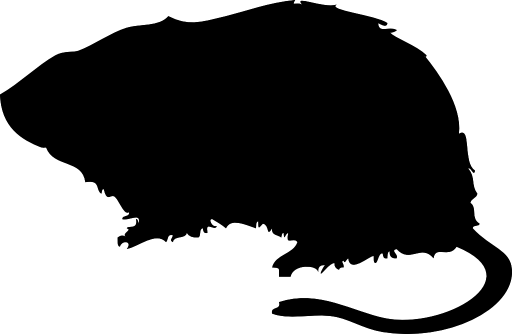 | Birds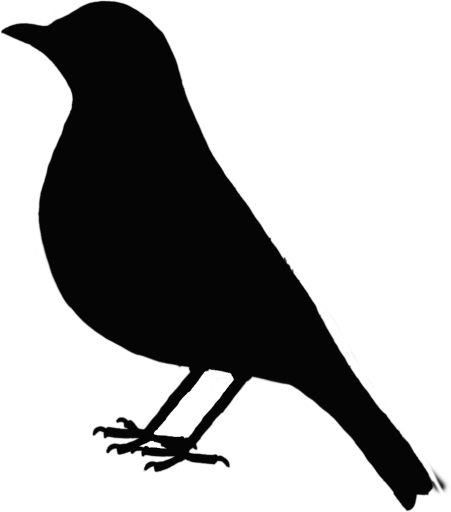 |
| --- | --- | --- | --- | --- |
| Taxonomic | Abiotic + biotic | 42 | 28 | 39 |
|  | Abiotic | 33 | 19 | 24 |
|  | Biotic | 28 | 16 | 33 |
|  | Residual | 10 | 9 | 17 |
| Phylogenetic | Abiotic + biotic | 92 | 68 | 85 |
|  | Abiotic | 37 | 64 | 40 |
|  | Biotic | 78 | 16 | 77 |
|  | Residual | 64 | 5 | 48 |
| Functional | Abiotic + biotic | 51 | 35 | 26 |
|  | Abiotic | 39 | 17 | 24 |
|  | Biotic | 30 | 30 | 2 |
|  | Residual | 16 | 13 | 2 |

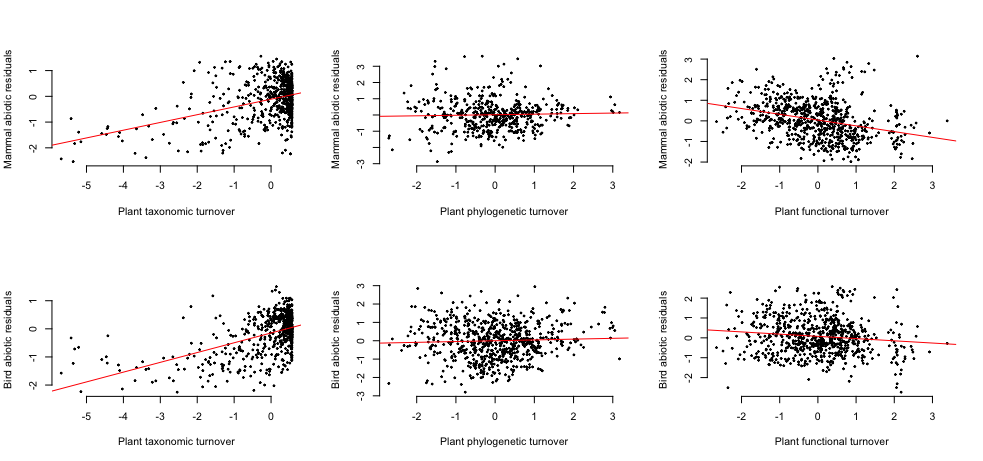

Appendix S6. Residuals from mammal and bird abiotic turnover models as a function of plant turnover. For each diversity metric, including taxonomic diversity (Sorensen's Index, left), phylogenetic diversity (PCDp, middle panels), and functional diversity (Functional Dissimilarity, right), residuals from models using only abiotic explanatory variables (environmental PC1, PC2, and elevation) are plotted as a function of plant diversity.
